# Four-Core Genotypes mice harbour a 3.2MB X-Y translocation that perturbs Tlr7 dosage

**DOI:** 10.1101/2023.12.04.569933

**Authors:** Jasper Panten, Stefania Del Prete, James P. Cleland, Lauren M. Saunders, Job van Riet, Anja Schneider, Paul Ginno, Nina Schneider, Marie-Luise Koch, Moritz Gerstung, Oliver Stegle, James M. A. Turner, Edith Heard, Duncan T. Odom

## Abstract

The Four Core Genotypes (FCG) is a mouse model system heavily used to disentangle the function of sex chromosomes and hormones. We report that a copy of a 3.2 MB region of the X chromosome has translocated to the Y*^Sry-^* chromosome and thus increased the expression of multiple genes including the auto-immune master regulator *Tlr7*. This previously-unreported X-Y translocation complicates the interpretation of studies reliant on FCG mice.

## Main text

Sex-dependence of mammalian physiology and disease is shaped by both gonadal hormones (testicular vs ovarian) and sex chromosome complement (XX, XY or variations thereof). The “Four Core Genotypes” (FCG) mouse model was developed to decouple the relative contributions of sex hormones and chromosomes by moving the testis determinant *Sry* from the Y chromosome to an autosome^1^. The FCG model thus generates mice that are: XX with ovaries (XXO), XX with testes (XXT), XY with ovaries (XYO), XY with testes (XYT) (Figure 1a). The FCG mouse model has been used in at least 90 primary research articles (Supplementary Table 1)^2^, affording insights into sex-biased immunity, cancer, Alzheimer’s occurrence and obesity^3–7^.

**Figure 1.**
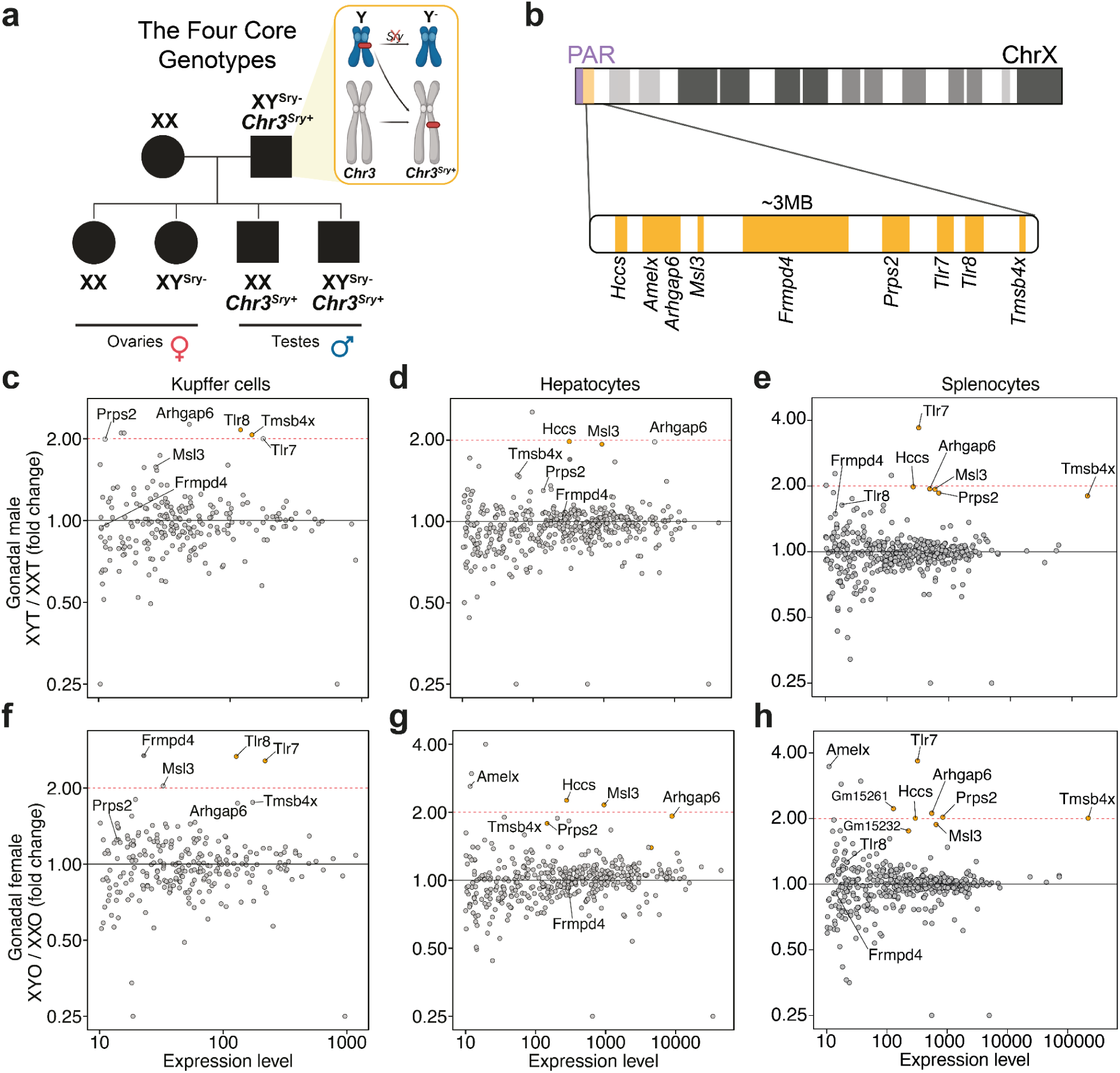
Transcriptional dosage of genes within a PAR-adjacent 3.2 MB region of the X chromosome is doubled in FCG XY mice. **a**, Schematic representation of the FCG model. **b,** A 3.2MB region flanking the PAR of X chromosome, containing nine genes. **c-d,** Differential expression analysis of pseudo-bulked single-cell RNA sequencing from all spleen cells between (c) XYT and XXT mice and between (d) XYO and XXO mice. **e-f,** Differential expression analysis of pseudo-bulked single-nucleus RNA sequencing of liver (e) Kupffer cells and (f) hepatocytes isolated from XYT and XXT mice. The x-axis indicates the expression level as total read counts. Highlighted in orange are all genes in the PAR-adjacent region that meet the expression level threshold. Gm15261 and Gm15232 are non-coding transcripts located in the same region.

The popularity of the FCG model has recently surged in parallel with the increased attention on sex-biased and hormone-driven differences in health and disease. Here, we report that this model has a previously-unreported copy of a multi-megabase region of the X chromosome fused to the Y chromosome with the *Sry* deletion (denoted Y*^Sry-^*), thus substantially increasing the expression level of several X-linked genes including *Tlr7* and *Msl3*.

While analysing single-cell RNAseq data from the spleen and liver of adult FCG mice (Supplementary Figures 1-4), we noticed that several genes located on a single contiguous region of the X chromosome adjacent to the pseudoautosomal region (PAR) (Figure 1b) are approximately 2-fold upregulated in XYT and XYO cells compared with their XX counterparts (Figure 1c-h, Supplementary Figure 5-6). This observation was unexpected, because X-inactivation should ensure equal X-linked gene dosage between XX and XY genotypes. We therefore hypothesised that a segment of X has translocated to the Y*^Sry-^*. Fusions with other chromosomes are unlikely to have survived the extensive backcrossing used to move the FCG model into a C57BL/6J (hereafter referred to as B6) background^8^. The identified set of nine X-linked genes showed cell type-specificity in their level of transcription. For example, immune cells in both organs consistently showed pronounced *Tlr7* expression bias towards the XY genotype. Further information on the cell type specificity of expression can be found in Supplementary Figure 7.

We then directly identified the amplified region via whole genome sequencing of the two FCG founder B6 XYT mice obtained commercially, alongside a wild-type B6 male with paired-end short-read sequencing (Figure 2a,b Supplementary Figure 8-10). This revealed that the amplification is a clean duplication of B6 sequence spanning a 3.2 MB, adjacent to the PAR of the X chromosome. We further resolved the bounds of the amplified region by whole-genome sequencing of FCG XYT and wild-type male mice on an F1 B6-FCG x CAST/EiJ (CAST) background, where the duplicated X-linked region can be genetically identified as originating from both the maternal CAST X chromosome and the paternal B6 genome (Figure 2c, Supplementary Figure 11). This demonstrates that the duplication is paternally inherited and likely fused to the Y chromosome. In total, we identified the amplification as approximately a 3.220.000 base pair region mapping to chrX:165.530.000 – 168.750.000 on *GRCm39*, containing nine annotated genes (*Tmsb4x, Tlr8, Tlr7, Prps2, Frmpd4, Msl3, Arhgap6, Amelx and Hccs*), of which six are detectably upregulated in an FCG XY-biased manner in the cells we profiled. To directly test whether the identified X amplification has translocated to the Y, we performed DNA FISH detection of both the whole Y chromosome and a ∼200kb domain located in the middle of the X chromosome amplification in wild-type male and FCG XYT primary splenocytes (Figure 2d). We observe that >90% of FCG XYT cells harbour two X domains instead of one (Figure 2e) and that one of these two X domains is in close proximity to the Y chromosome (Figure 2f). Thus, multiple lines of evidence support that the increased expression observed in FCG XY genotype mice is caused by an X-Y translocation.

**Figure 2.**
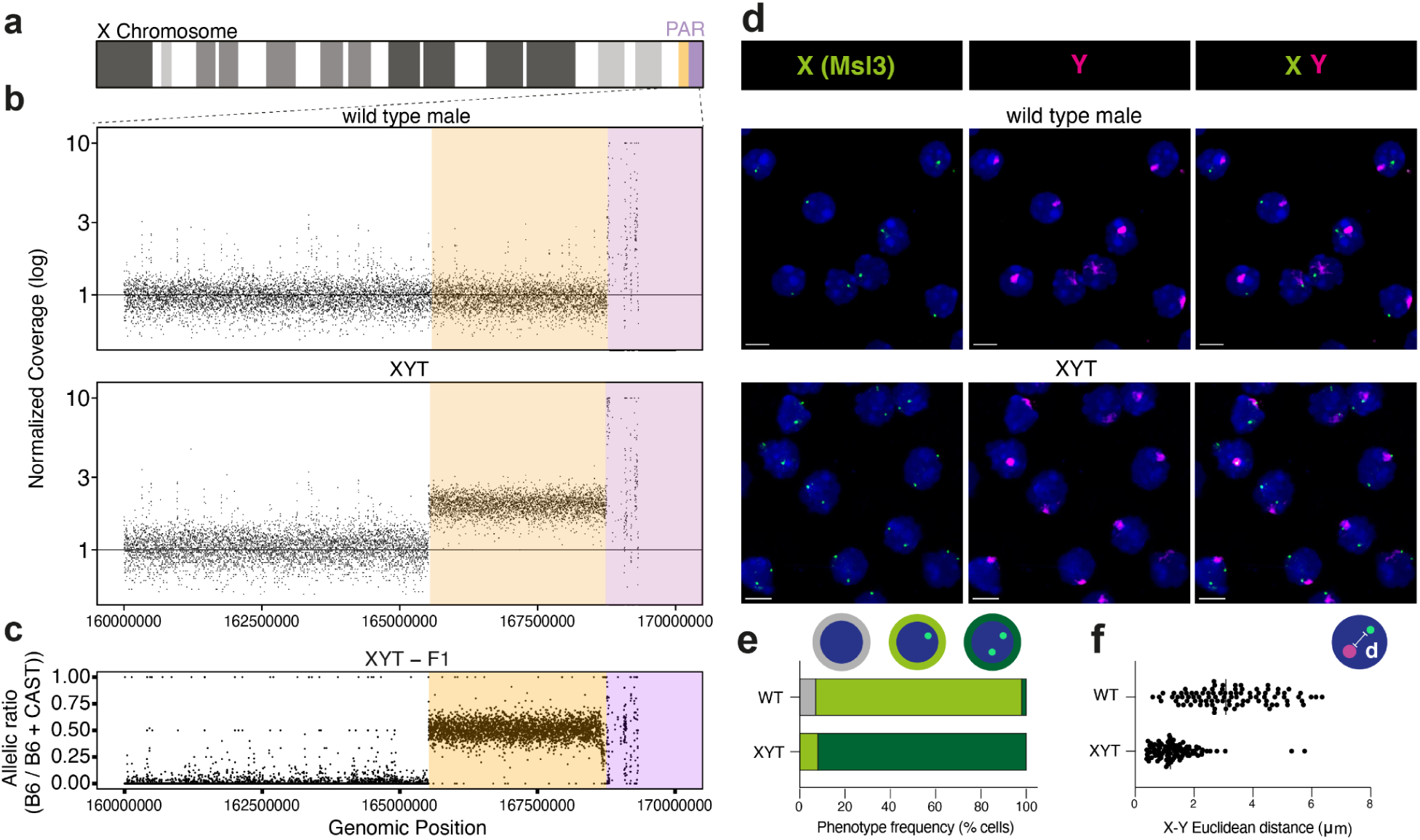
Whole genome sequencing and DNA FISH reveal an X-Y translocation in FCG XY mice. **a,** Overview of the X-chromosome, where orange indicates the location of the amplified region, and purple indicates the PAR. **b,** Coverage of sequencing reads normalized to the median autosomal coverage in 1kb windows of a C57B6 wild-type male (top) and the FCG XYT founder mouse (bottom). **c,** Ratio of haplotype-resolved coverages in an FCG XYT F1 hybrid. The presence of B6-specific reads in the amplified region reveals a paternally inherited 3.2 MB region duplicated in the FCG model. **d.** DNA FISH of the whole Y chromosome (magenta) and a 200kb domain of the X amplification (green) in C57B6 wild-type male (top row) and FCG XYT (bottom row) primary splenocytes. Images represent maximum intensity projections. Scale bars represent 5µm. **e-f.** Quantification of X domain number (e) and Euclidean distance between the Y domain and its nearest X domain (f). n = 100 cells across five fields of view per genotype. Error bars represent standard deviation (f).

Our discovery that a copied 3.2 MB segment of the X has become fused to the Y*^Sry-^* chromosome reflects the long-known proclivity for regions adjacent to the PAR to undergo genetic exchange^9^. Indeed, a similar translocation has occurred in the YAA model, wherein a 4 MB stretch of the X chromosome is fused to the Y chromosome, with consequences to gene expression and susceptibility to auto-immunity disease^10^.

While we purchased our XYT colony founders in summer 2022, the model has been commercially available following extensive backcrossing to move the FCG model onto a C57BL/6J background, completed in 2009^8^. Upregulation of the translocation genes we identified in FCG XY mice has been previously reported^11–14^, suggesting that this genetic alteration has been circulating for at least a decade.

The increased dosage of these nine duplicated genes could impact prior studies and any ongoing projects that rely on the FCG model. The most obvious recommendation is that genetic models perturbing sex chromosome complement should be routinely genome-sequenced, due to inherent genetic instability of PAR-adjacent regions^9,10,15^. Even the FCG mouse model carrying an X-Y translocation can be reliably used to identify gonadal hormone contributions in the XXO and XXT groups. Our findings emphasize the importance of cross-validating results arising from the FCG model using other sex chromosome complement models and wild-type males, as has long been recommended^7^.

## Methods

### Mouse colony management

Four Core Genotypes colony founders on the C57BL/6J background were obtained from The Jackson Laboratory (strain 010905)^8^. Two XYT individuals were mated with wild-type C57BL/6J (Janvier-Labs) or CAST/EiJ (bred at the German Cancer Research Center) females to respectively generate pure C57BL/6J and B6 x CAST F1 hybrid XXO, XXT, XYO and XYT progeny. Wild-type C57BL/6J males (Janvier-Labs) were imported and housed for one week prior to sacrifice. All animals were maintained as virgins (apart from the two XYT founders) and housed in groups of up to six mice in Tecniplast GM500 IVC cages with a 12 hour light/dark cycle. Mice had ad libitum access to water, food (Kliba 3437) and environmental enrichment. The colony was regularly controlled for infections using sentinel mice to ensure a healthy status. All breedings and organ collections were carried out with the approval of the Regierungspräsidium Karlsruhe. Animal genotype (i.e. XXO, XXT, XYO or XYT) was assigned based on an independent assessment of sex chromosome complement and gonadal sex. Sex chromosome complement was initially determined by PCR detection of the Y chromosome using published primers YMTFP1 (5’ CTG GAG CTC TAC AGT GAT GA 3’) and YMTRP1 (5’ CAG TTA CCA ATC AAC ACA TCA 3’)^16^ and confirmed by detection of *Xist* (specific to the XX genotypes) and *Sry* (specific to the XY genotypes) sequencing signal. Gonadal sex was determined by visual assessment of external morphology, initially at the time of weaning and confirmed at the time of sacrifice.

### Tissue collection and processing

Animals were sacrificed by carbon dioxide inhalation at 12 weeks of age unless otherwise stated. Whole livers were dissected out, transferred to 5ml tubes, snap-frozen on liquid nitrogen and stored at –80°C. Frozen livers were pulverised with the cryoPREP® Automated Dry Pulverizer (Covaris #CP02) using the TT2 holder, TT2 consumables and impact setting 6 and pulverised material stored at –80°C. Whole spleens were dissected, transferred to a petri dish containing DMEM (Gibco # 41966029) with 5% fetal bovine serum (FBS, Thermo Fisher # 16140071) and processed directly for single cell isolation. All tissue collections were performed between 8:00 and 12:00 to approximately control for circadian regulation of organ physiology.

### Single-cell RNA sequencing: sample preparation, library preparation and sequencing

Dissected spleen fragments were transferred to a 40-μm cell strainer (Greiner # 542040) and ground with a syringe plunger. The cell suspension was strained twice. Strainers were washed with additional DMEM (Gibco # 41966029) with 5% FBS and cells were pelleted by centrifugation at 300 x g for 4 minutes at 4°C. Cell pellets were carefully resuspended in 300 uL of ACK lysis buffer (Lonza # BP10-548E) for 1 minute in ice. After incubation cells were washed with 1mL of DMEM with 5% FBS and pelleted. Cells concentration was determined using 0.4% trypan blue stain (Invitrogen) and Countess II automated cell counter (Invitrogen). 1 × 10^7 cells were processed for dead cell depletion using Dead Cell Removal Kit (Miltenyi Biotec # 130-090-101) as per the manufacturer’s instructions. The magnetically labelled dead cells were retained using MS MACS Columns (Miltenyi Biotec # 130-042-201) on a MiniMACS Separator (Miltenyi Biotec # 130-042-102). The live cell fraction was pelleted and resuspended in 1X PBS with 0.04% bovine serum albumin (BSA, Miltenyi Biotec # 130-091-376) to a final concentration of 1 × 10^6^ cells/mL (1,000 cells per μL).

Single-cell RNA was performed with 20.000 cells which were encapsulated into droplets in the Chromium Controller instrument using Chromium Next GEM Single Cell 3’ Reagents Kits v3.1, according to the manufacturer’s recommendations. Briefly, single cell lysis and barcoded reverse transcription (RT) of mRNA happened into the droplets. Twelve cycles were used for cDNA amplification and sample index PCR to generate Illumina libraries. Quantification of the libraries was carried out using the Qubit dsDNA HS Assay Kit (Life Technologies), and cDNA integrity was assessed using D1000 ScreenTapes (Agilent Technologies). Libraries were pooled in an equimolar manner and 1000pM were paired-end sequenced on an Illumina NextSeq2000 (R1 = 30 cycles, R2 = 8 cycles, I1 = 0 cycles, I2 = 199 cycles).

### Single-cell RNA sequencing: data processing and analysis

Genomic references for C57BL/6 (GRCm38) was generated using CellRanger *mkref* (v7.0) using sequence and gene annotations from Ensembl (release 94). For mapping of C57B6 / CAST/EiJ F1 mouse data, a joint reference was constructed based on the GRCm38 reference in which SNP positions between mm10 and CAST/EiJ were N-masked. These SNPs were derived from (Keane2011, ftp://ftp-mouse.sanger.ac.uk/current_snps/mgp.v5.merged.snps_all.dbSNP142.vcf.gz). Filtered count matrices were generated using CellRanger *count* (v6.1.1) using default settings, which includes reads mapped to introns. Low-level analysis of scRNA-Seq data was then performed, largely using functions from the *scran* (v1.24.1) and *scater* (v1.20.1) R packages^17,18^. First, cells with less than 500 UMIs and 500 detected genes were removed. Next, counts were normalised using the *computeSumFactors* function and log-transformed. For cell type annotation, we used mutual nearest neighbour-based batch correction using the function *MNNcorrect* with the library as batch variable to exclude species-specific and technical variation across samples^19^. The resulting corrected matrix was used for dimensionality reduction by principal component analysis (*prcomp, stats*), tSNE (*Rtsne, Rtsne,* v0.15) and UMAP (*umap, umap,* v0.2.7.0). To identify cell type clusters, we used graph-based community detection using the Louvain algorithm implemented by the functions *buildSNNGraph* and *cluster_louvain* of the package *igraph* (v1.2.10). Cell type labels were defined by label transfer from the ImmGenData dataset (celldex R package, v1.6.0) using the SingleR R package (v1.10.0). Doublets were identified and excluded using the scDoubletFinder package (v1.15.4).

### Single-nucleus RNA sequencing: sample preparation, library preparation and sequencing

Nuclei isolation was performed as described previously^20^, with the following modifications. ∼20mg pulverised frozen liver tissue was dounced ten times with pestle A and ten times with pestle B (Sigma # 8938) in 1ml hypotonic lysis buffer solution B containing 10 mg/ml BSA (Sigma # 126609). The homogenate was then diluted 1:5 in fresh lysis buffer, incubated on ice for 10min, filtered through a 30µm CellTrics filter (Sysmex # 04-004-2326) and spun at 700 rcf for 5min. Pelleted nuclei were resuspended in 1ml SPBSTM supplemented with 1% DEPC, spun again and resuspended in 1ml SPBSTM. Nuclei were fixed by addition of 4.3ml pre-chilled fixation solution (4ml MeOH and 300ul 16.7mg/ml DTSSP, Sigma # 803200) and incubation on ice for 15min, rehydrated by addition of 3 volumes of SPBSTM, spun at 700 rcf for 10 min, resuspended in 1ml SPBSTM and stored at –80 in 250 µl aliquots. Nuclei quality and concentration was quantified using a LUNA-FX7 automated cell counter (Logos). Library preparation was performed as described previously^20^, with the following modifications. Nuclei were passed through a 20µm EasyStrainer filter (Greiner # 542120) prior to distribution across the reverse transcription plate. Following protease digestion, the optimal Tn5 concentration was empirically determined by testing a 2-fold dilution series (0.25, 0.13, 0.06, 0.03 N7-loaded Tn5 per 5ul reaction) on a small subset of wells (2 per dilution). Final libraries were subjected to a two-sided (0.45×/0.80×) AmpureXP bead cleanup but not gel extraction. Libraries were sequenced on a NovaSeq 6000 system (Illumina) with a S4 200 cycles reagent kit (R1 = 34 cycles, R2 = 84 cycles, I1 = 10 cycles, I2 = 10 cycles).

### Single-nucleus RNA sequencing: data processing and analysis

#### Sample demultiplexing and alignment

For handling the demultiplexing and subsequent downstream alignment and read-counting of sci-RNA-seq3 data, we designed a custom *Snakemake*^21^ workflow termed sci-rocket which has been made publicly-available under the MIT licence. (https://github.com/odomlab2/sci-rocket). Raw sequencing data (binary base calls) were converted to paired-end read sequences using *bcl2fastq* (v2.20.0.422) with the PCR index #1 and PCR index #2 sequences added within the read-name using the following parameters:

`bcl2fastq –R <bcl> –-sample-sheet <sample_sheet> –-output-dir <out> –-loading-threads <n> –-processing-threads`

`<N> –-writing-threads <N> –-barcode-mismatches 1 –-ignore-missing-positions –-ignore-missing-controls,`

`--ignore-missing-filter –-ignore-missing-bcls –-no-lane-splitting –-minimum-trimmed-read-length 15`

`--mask-short-adapter-reads 15`

Subsequently, raw paired-end sequences were split into smaller evenly-sized chunks (*n*=75) using *fastqsplitter* (v1.2.0) which were analysed in parallel during the subsequent sample-demultiplexing procedure. Reads were assigned to their respective sample based on the combinatorial presence of four barcodes within the R1 read; PCR index #1 (p5), PCR index #2 (p7), ligation primer and the reverse transcriptase (RT) primer used during the sci-RNA-seq3 protocol. Each barcode was 10 nucleotides in length, except for the ligation barcode which could be either 9 or 10 nucleotides in length. From each R1 read, these barcodes were retrieved and matched, with a hamming-distance of max. 1nt, to the white-list of barcode sequences which could be present within the sequencing run. If all four barcodes could be matched, each paired-end read was deposited into their respective sample-specific .fastq files. The read containing the combinatorial barcode (R1) was modified to only contain the sequence of the matching white-listed barcodes (if hamming distance > 0) and UMI of 8nt in length. Ligation barcodes of 9 nucleotides in length were padded with an extra G in order to always produce sequences of 48 nucleotides in length; PCR Index #1 (10nt), PCR index #2 (10nt), ligation (10nt), RT (10nt) and UMI (8nt). The ligation white-list was also adjusted accordingly. Read-pairs without all four barcodes matching the white-lists were discarded into a separate file containing all discarded reads with information on which barcode sequence(s) were non-matching. Demultiplexed .fastq files from each parallel job (*n*=75) were merged to produce a single sample-specific paired-end .fastq file (R1 and R2).

After sample-demultiplexing, remaining adapter sequences and low-quality bases (≥Q15) were trimmed using *fastp* (v0.23.4)^22^. Read-pairs with mates containing fewer than 10 nucleotides after trimming were discarded. Alignment of trimmed reads was performed against GRCm39 (M31) with GENCODE (v31) annotations^23^ using STARSolo (v2.7.10b)^24^ with the following parameters:

`STAR –-genomeDir <index> –-runThreadN <n> –-readFilesIn <R2> <R1> –-readFilesCommand zcat—soloType`

`CB_UMI_Complex –-soloCBmatchWLtype Exact –-soloCBposition 0_0_0_9 0_10_0_19 0_20_0_29 0_30_0_39`

`--soloUMIposition 0_40_0_47 –-soloCBwhitelist <wl_p7> <wl_p5> <wl_lig< <wt_rt> –-soloCellFilter CellRanger2.2`

`<n_expected_cells> 0.99 10 –-soloFeatures GeneFull –-soloCellReadStats Standard –-soloMultiMappers EM`

`--outSAMtype BAM SortedByCoordinate –-outSAMunmapped Within –-outFileNamePrefix <sample> –-outSAMmultNmax 1`

`--outSAMstrandField intronMotif –-outFilterScoreMinOverLread 0.33 –-outFilterMatchNminOverLread 0.33`

`--outSAMattributes NH HI AS nM NM MD jM jI MC ch XS CR UR GX GN sM`

In addition, indexes of BAM files were generated using sambamba (v1.0)^25^. Briefly, this generated sparse matrices containing the UMI counts for each gene per cellular barcode. For each gene, we counted the total set of UMI of all intronic, exonic and UTR-overlapping reads based on GENCODE (v31) annotation. Reads mapping to multiple genes (multi-mappers) were counted using the Expectation-Maximization (EM) algorithm available within STARSolo. After visual inspection of the sample-wise knee-plots with the STARSolo-designated UMI threshold to distinguish ambient RNA from true cell, we opted to adopt these UMI-thresholds as-is.

#### Data import and quality control

Using monocle3 (v1.3.1)^26^, we combined the sample-wise UMI matrices generated by STARSolo and imported all protein-coding, lncRNA and immunoglobulin genes and removed predicted-only genes for a total of 25,064 genes across 131,975 cells. Next, we determined the number of relevant principal components capturing at least ≥1% of variance (*n*=20) and used these principal components in subsequent dimension reduction using the default monocle3 workflow whilst using the number of UMI per cell to regress out potential batch-effects. Initial major cell-type clusters were identified by Leiden clustering (*k*=20, resolution=1^e-6^)^27^. To detect potential doublets, we performed scDblFinder (v1.14.0)^28^ to estimate captured cells which resembled artificial combination(s) of multiple major clusters (*n*=16) within our experiment. We utilised all relevant principal components capturing at least ≥1% of variance (*n*=20), assumed a maximum doublet rate of max. 2% (*dbr*=0.02) and used the default number of artificially-generated doublets. In addition, cells with a total UMI count ≥ µ * 3σ (based on all cells) were flagged as being potential outlier**s (*n*=**4,526). After visual inspection of cells flagged as doublet or UMI-outlier **(n=**4,626; 1.1% of total cells) within UMAP-space, we removed these cells prior to downstream analysis.

#### Differential expression testing

Differential expression analysis of single-cell and single-nucleus RNA-Seq data was performed as a pseud-bulk per individual mouse and using the DESeq2 package^29^. To this end, raw read counts for each library were summed up, and genes with fewer than 10 reads per sample were excluded as lowly expressed. Then DESeq2 (v1.36.0) was used to compute size factors (*estimateSizeFactors* function) and to detect differentially expressed genes between XY and XX genotypes with default parameters (*DESeq* function), separately for the two gonadal groups. Genes were considered differentially expressed at an FDR of 10% (Benjamini-Hochberg correction). To decrease the multiple testing burden, we only test genes on the X-chromosome in this analysis. The single-nucleus RNA-Seq dataset was analyzed in the same way, but separating individual cell types.

#### Whole Genome Sequencing

DNA was isolated from ∼30 mg frozen liver and 2mm^3^ ear punch biopsies using a protocol incorporating elements of the AllPrep DNA/RNA Micro Kit (Qiagen) and a DNeasy Blood and Tissue Kit (Qiagen). Frozen tissue samples were homogenised by adding 600 uL RLT containing 1% β-Mercaptoethanol and a 5mm stainless steel bead (Qiagen # 69989) and processing for 40 seconds at 15 Hz with the TissueLyser II. Lysed samples were supplemented with 20ul proteinase K (from the Blood and Tissue Kit) and incubated at 56°C for 10 min. Half of the digested sample was combined with 500uL Buffer AL (Qiagen Blood & Tissue), mixed by vortexing and incubated at 56°C and 500 rpm for 30 min. An equal volume of absolute ethanol was added to the samples and mixed thoroughly. The precipitated mixture was transferred to a DNeasy Mini column and washed twice with 500 μl buffer AW1 and subsequently with 500 μl AW2. After drying the membrane for 2 minutes at 20,000 x g, DNA was eluted using 50 μl AE buffer pre-warmed to 70°C. DNA quality and quantity were assessed using a Genomic DNA ScreenTape (Agilent Technologies) and Qubit dsDNA BR Assay Kit (Life Technologies).

Dual Index whole genome libraries were prepared with 300ng of genomic DNA using the NEBNext Ultra II kit (E7805S/L). Fragmentation time was optimised and occurred at 37°C for 20min. Adaptor-ligated DNA was size selected with AMPure XP Beads (Beckman Coulter) to select an insert size of approximately 150-200bp. Libraries were amplified 6 cycles using the NEBNext UDI primers (E6440). Library size and molarity were determined using a D5000 ScreenTape (Agilent Technologies) and Qubit dsDNA HS Assay Kit (Life Technologies). Final library size was approximately 600-700bp. Libraries were pooled in an equimolar manner and 750pM was sequenced, paired-end 150bp, using the NextSeq 2000 platforms (300 cycle P3 chemistry) to aim for 200Mio reads per sample.

#### Whole-Genome Sequencing: data processing and analysis

For the whole genome sequencing data, reads were first trimmed to using Trim Galore and cutadapt (https://github.com/FelixKrueger/TrimGalore) using the options `trim_galore –paired –-three_prime_clip_R1 1 –-three_prime_clip_R2 1 –-fastqc –-cores 8 –-stringency 3.` Next, reads were aligned to a GRCm39 in which all SNP positions between C57B6 and CAST/EiJ strains are N-masked (see also Single-cell RNA sequencing: data processing and analysis) using bowtie2 with default options ^30^. PCR duplicates were deduplicated when they mapped to the exact same positions using the samtools programs collate, fixmate, sort and markup. Finally, for the F1 mouse samples, reads mapping to the C57B6 or CAST/EiJ genomes were assigned to either parental haplotype (or unassigned) using SNPsplit^31^. From the sorted and deduplicated bam-files, both total and allele-specific coverage tracks with windowsizes 1kb and 1000kb were generated using the *bamProfile* function from the *bamsignals* package (v1.28.0).

#### DNA fluorescence in situ hybridization (FISH)

Primary splenocytes were isolated from five month old XYT-WT and XYT-FCG mice as described above for scRNA-seq. 10 million splenocytes were spun at 200 rcf and room temperature (RT) for 5 min, resuspended in 3:1 methanol:acetic acid fixative, incubated at –20°C for at least one hour, spun at 200 rcf and RT for 5 min, resuspended in 45% acetic acid, incubated at RT for 5min, spotted onto SuperFrost Plus Gold microscope slides, air dried overnight at RT and stored at 4°C until use. Labelled *Msl3* probes were generated using a nick translation kit (Abbott) and midiprepped bacterial artificial chromosome RP23-391N16 (BACPAC Genomics) as template. FISH was performed largely according to the instructions provided by the nick translation kit manufacturer (https://www.molecular.abbott/int/en/vysis-fish-knowledge-center/nick-translation-kit-preparing-the-reagents) with the following modifications. Cells were incubated in 0.1mg/ml RNase A and 10U/ml RNase H diluted in 2× SSC for 1h at 37°C^32^ prior to hybridization. Hybridisation was performed for 21h using 5ul Msl3 probes resuspended in Y chromosome paint solution (MetaSystems Probes #D-1421-050-OR) and 16mm diameter coverglasses. Samples were counterstained with 5 µg/ml Hoechst in 2× SSC for 5 min at RT, mounted with Prolong Diamond mounting medium and allowed to cure at RT for at least 24 h. Images (7µm Z-stacks with 1µm step size) were acquired with an IXplore Spin spinning disc confocal microscope equipped with a 100× oil immersion objective (Olympus). Five non-overlapping fields of view (FOV) were acquired per genotype. Images were quantified in Fiji using the Cell Counter plugin. For each genotype, 100 cells across five FOVs were counted. Specifically, cells with a clear Y domain were selected blind to X signal, then X domain number and X-Y distance (i.e. distance from the centre of the Y domain to its nearest X domain) were quantified.

### Data availability

The whole genome sequencing data has been submitted to ArrayExpress under the accession number E-MTAB-13585. The single-cell RNA-sequencing data has been submitted to ArrayExpress under the accession number E-MTAB-13586. ArrayExpress submission of the single-nucleus RNA-sequencing data is in progress and a revised version of this manuscript complete with the accession number will be uploaded as soon as the process is complete.

### Code availability

Our custom *Snakemake* workflow (sci-rocket) used in processing the sci-RNA-seq3 data has been made publicly-available under the MIT licence (https://github.com/odomlab2/sci-rocket). Additional custom code used for the processing, analysis and visualisation of data supporting this manuscript is available via github (https://github.com/jasperpanten/fourcore_transloc).

**Table 1:**
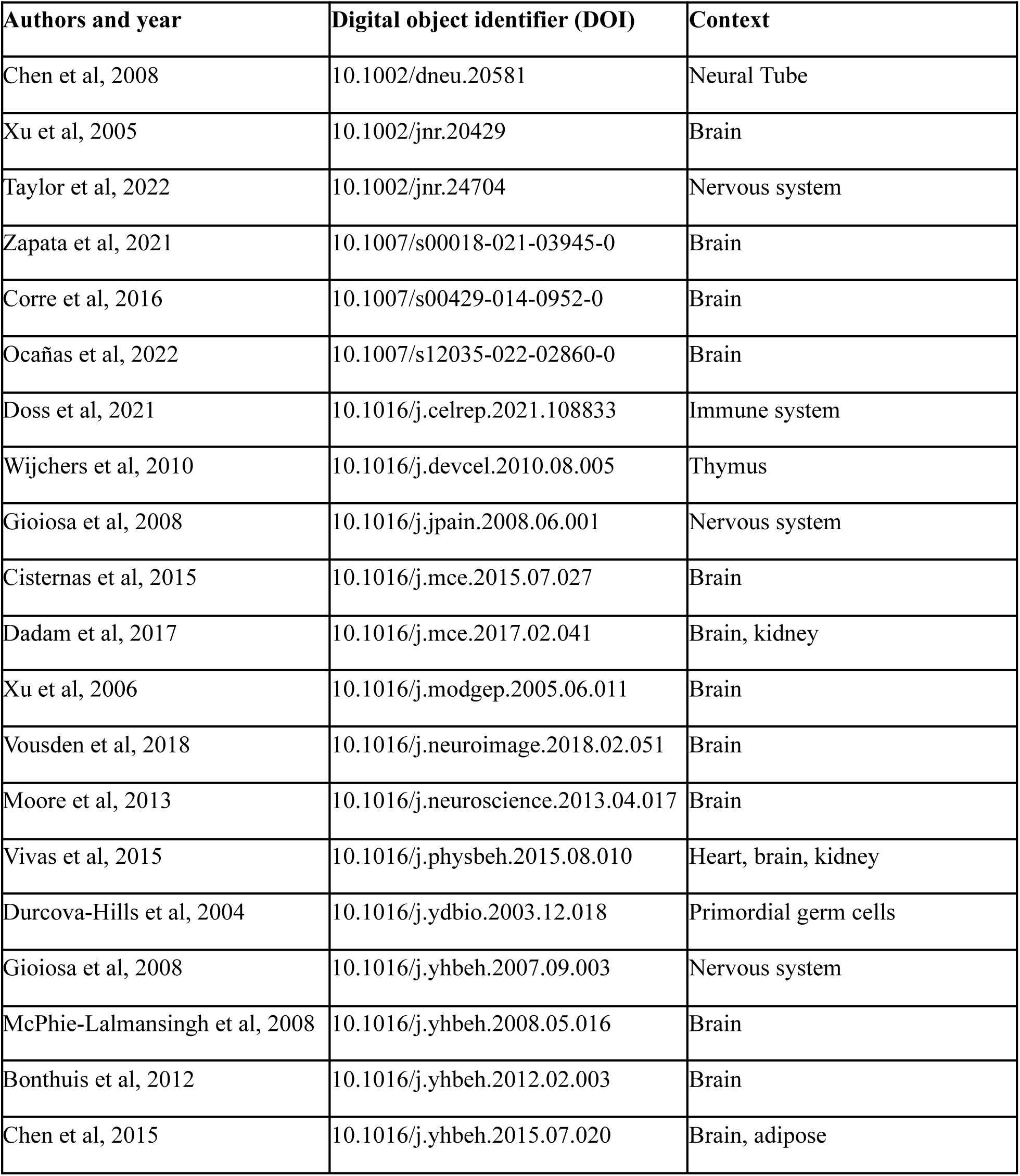
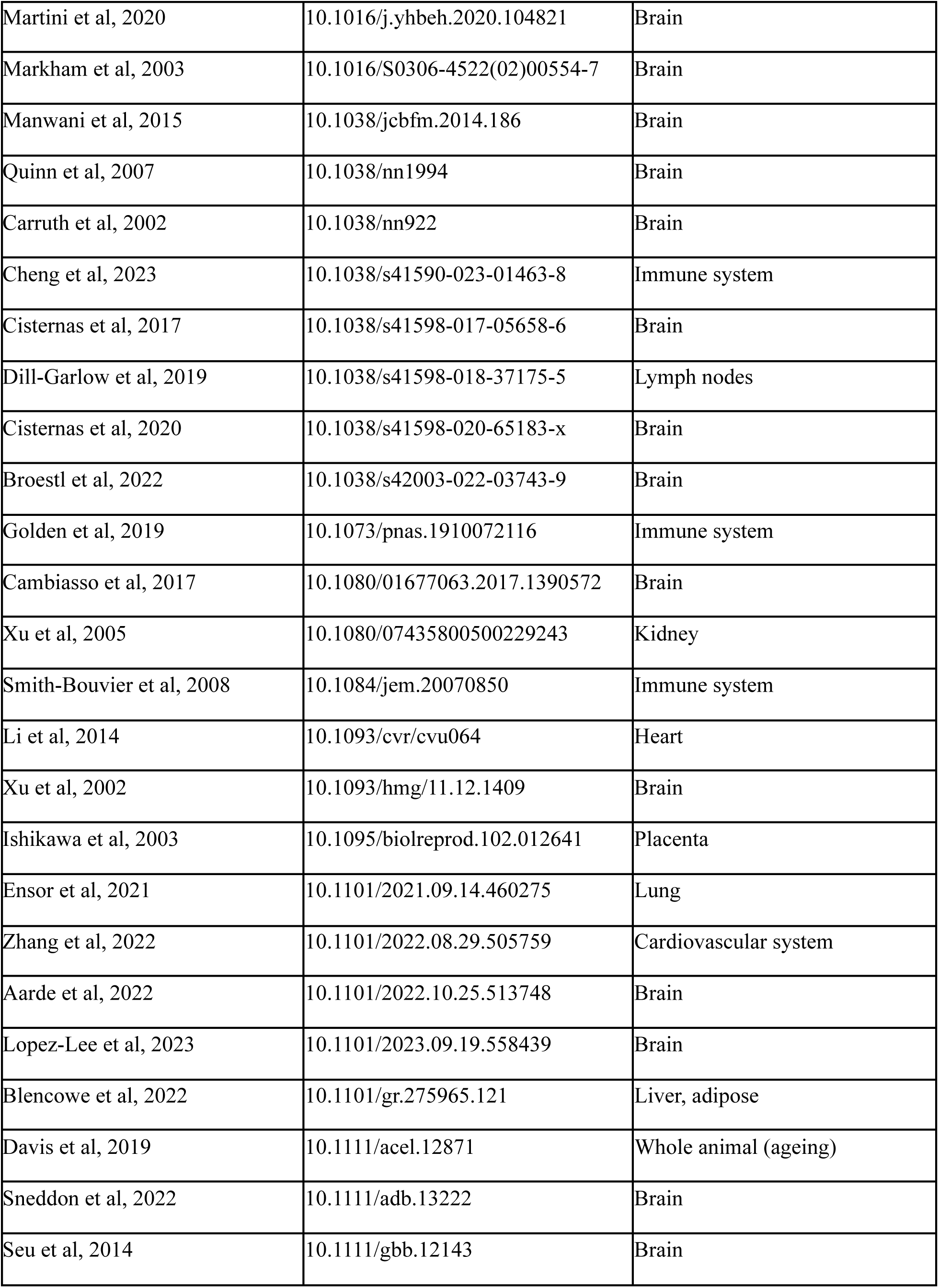
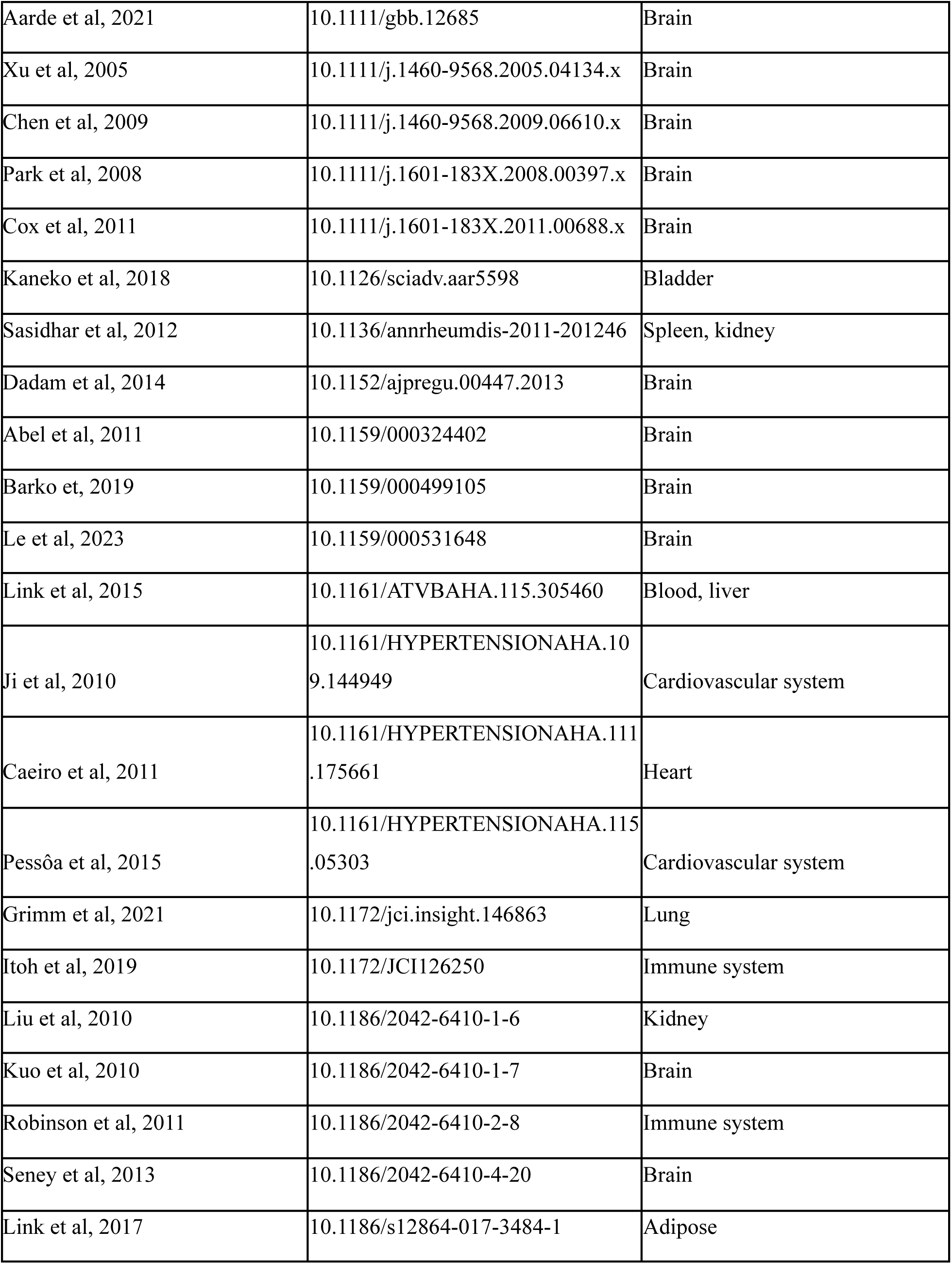
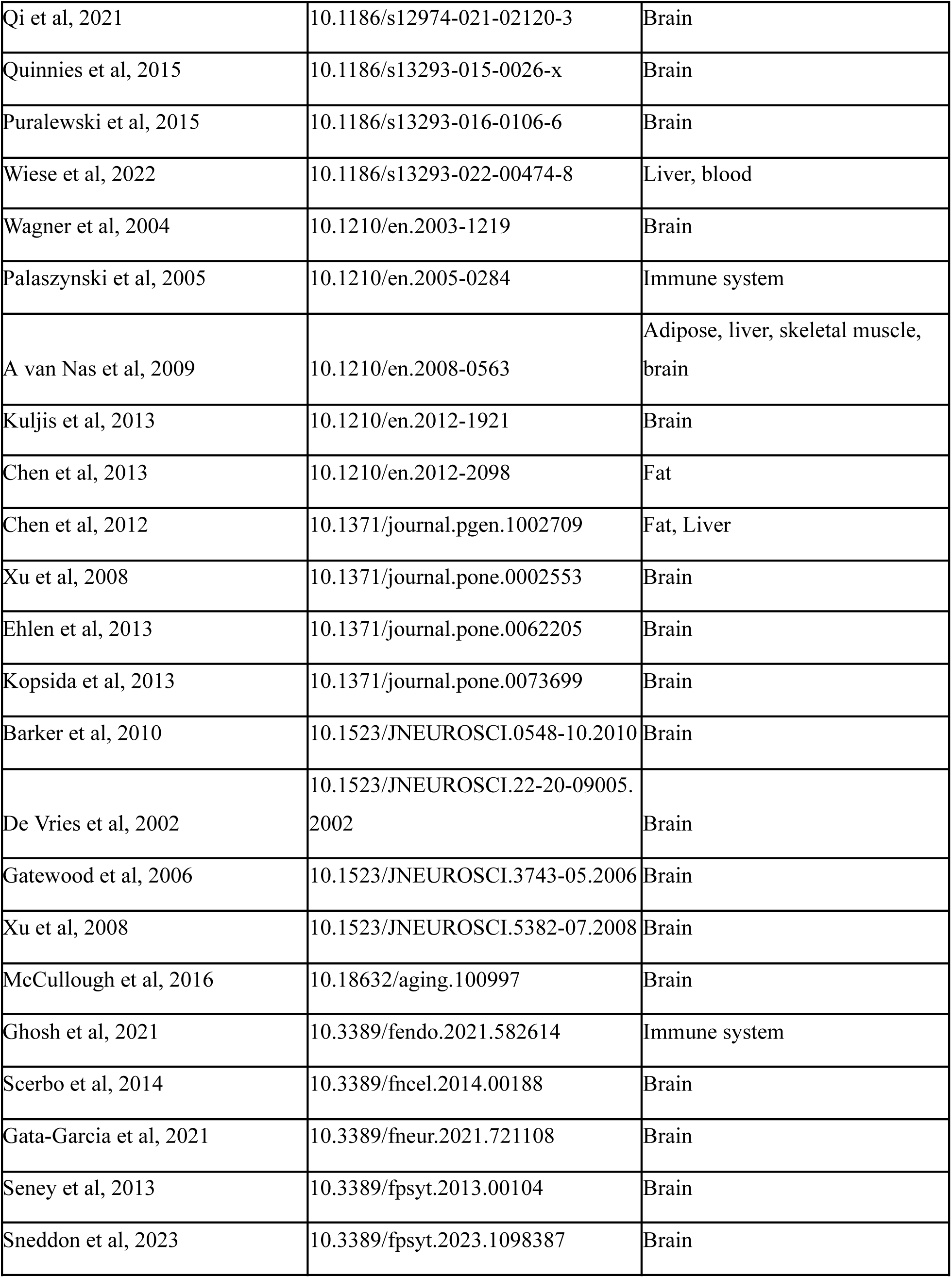
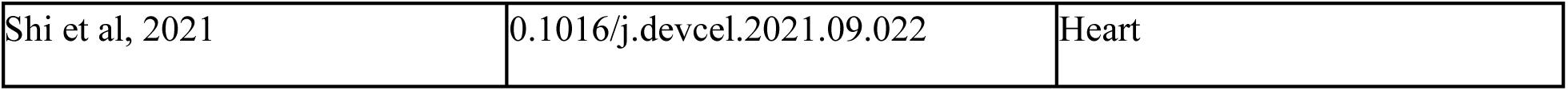
Publications using FCG mice. The following studies have made use of the FCG mouse model. We emphasise that the translocation does not necessarily affect the conclusions drawn in each of them, as many of them dissect phenotypes unaffected by the translocation and/or they validate results with orthogonal approaches, including other mouse models. Finally, the colonies used in these studies might not necessarily carry the translocation.

## Acknowledgements

We thank all members of the Odom and Heard labs (in particular Dr. Tim Pollex and Dr. Agnese Loda), the DKFZ Central Animal Laboratory and Next Generation Sequencing core facilities. We thank Prof. Arthur P. Arnold for discussions, manuscript feedback and tissue samples. This research was financially supported by core funding from the Helmholtz Association (to DTO) and EMBL (to EH), a grant from the European Research Council (788937 to DTO), a grant from the Wilhelm Sander Stiftung (to DTO and EH), a DKFZ Postdoctoral Fellowship (to JPC) and state funds approved by the State Parliament of Baden-Württemberg for the Innovation Campus Health + Life Science Alliance Heidelberg Mannheim (to JvR).

## Contributions

JP and SDP made the initial discovery. SDP performed single-cell RNA-Sequencing experiments with assistance from NS. JC and LS performed single-nucleus RNA sequencing with assistance from AS and MLK. SDP performed whole-genome sequencing with assistance from AS. JC performed FISH experiments. JP, SDP, LS and JvR performed computational analysis of sequencing data with assistance from PG. SDP, JC, AS and MLK managed the mouse colonies. JP, SDP, JC, LS and DTO generated figures. JT provided critical intellectual input. JP, SDP, JC and DTO wrote the manuscript with input from all co-authors. MG, OS, EH and DTO supervised the project. All authors had the opportunity to edit the manuscript. All authors approved the final manuscript.

## Supplementary Figures

**Supplementary Figure 1:**
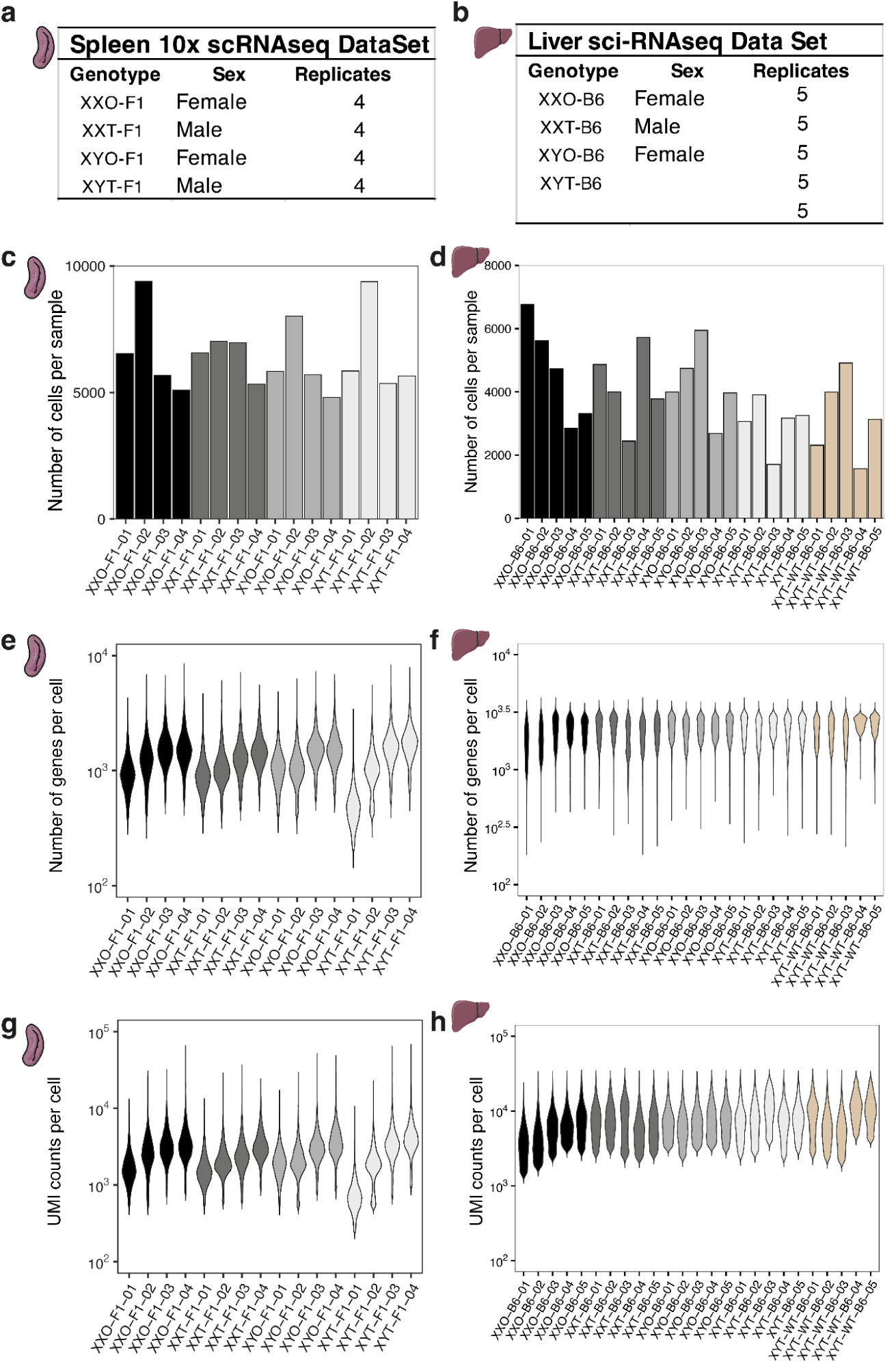
Quality control of the single cell and single nucleus datasets. **a-b**, Overview of samples used for transcriptome analysis**. c-d,** Number of cells recovered per biological replicate. **e-f,** Median number of genes per cell in each biological replicate. **g-h,** Median number of UMI (Unique Molecular Identifier) per cell in each biological replicate. The number of cells, genes and UMIs are consistent between the different replicates.

**Supplementary Figure 2:**
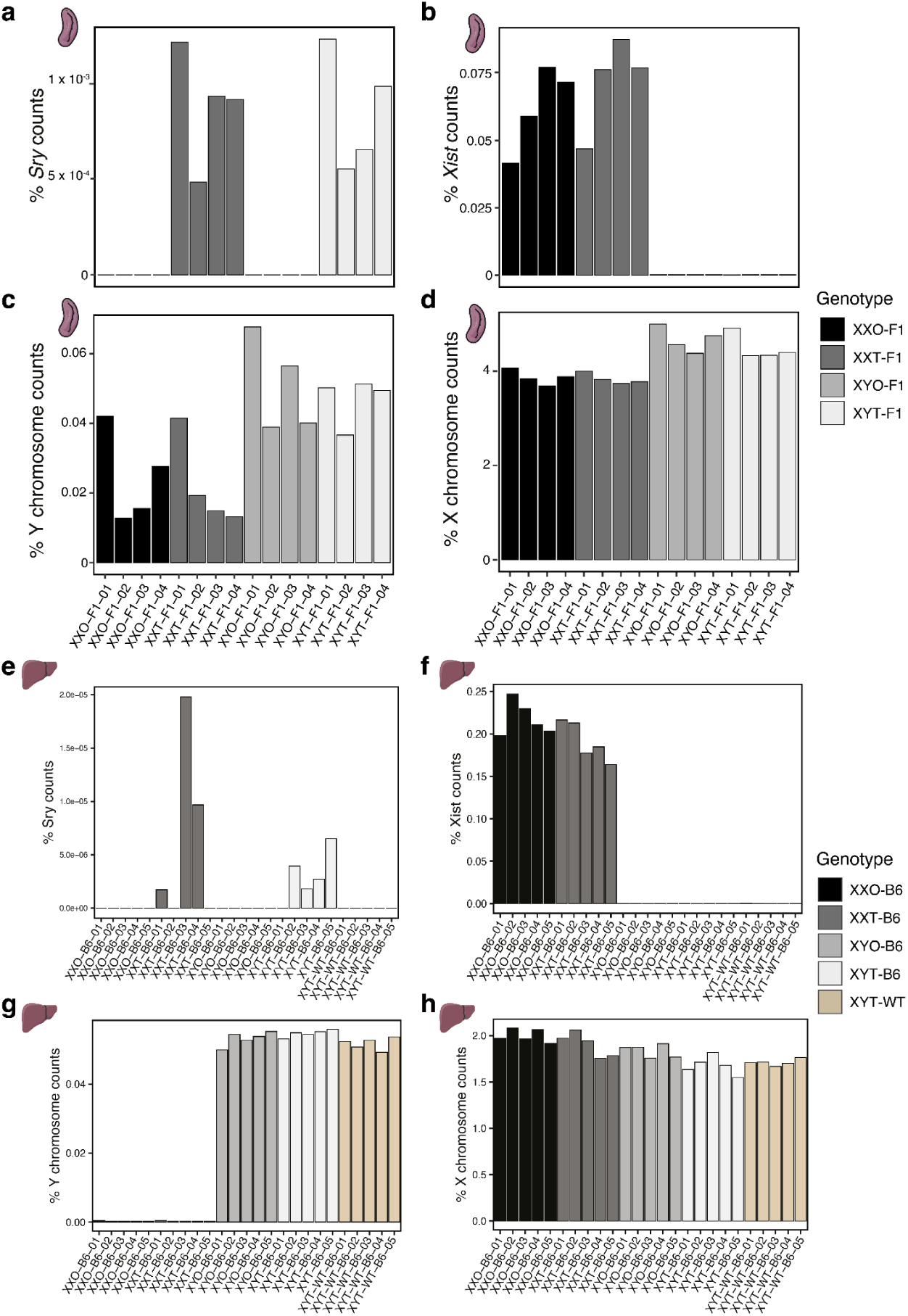
Genotyping of the mice by transcriptomic data. **a,e**, *Sry* expression in each biological replicate calculated as a percentage of Sry counts / Total counts. The data confirmed the presence of *Sry* in the FCG gonadal male mice (XYT and XXT). **b,f,** *Xist* expression in each biological replicate calculated as a percentage of Xist counts / Total counts. As expected, *Xist* is expressed only in mice carrying two X chromosomes. **c,g,** Y chromosome expression in each biological replicate calculated as a percentage of Y chr. counts / Total counts. Y chromosome counts are only present in the mice carrying a Y chromosome. **d,h,** X chromosome expression in each biological replicate calculated as a percentage of X chr. counts / Total counts. In spleen (**d**) the counts are higher in XY genotypes, likely because of the high expression of *Tmsb4x*. In liver (**h**), the read counts are comparable, showing dosage compensation through X-chromosome inactivation.

**Supplementary Figure 3:**
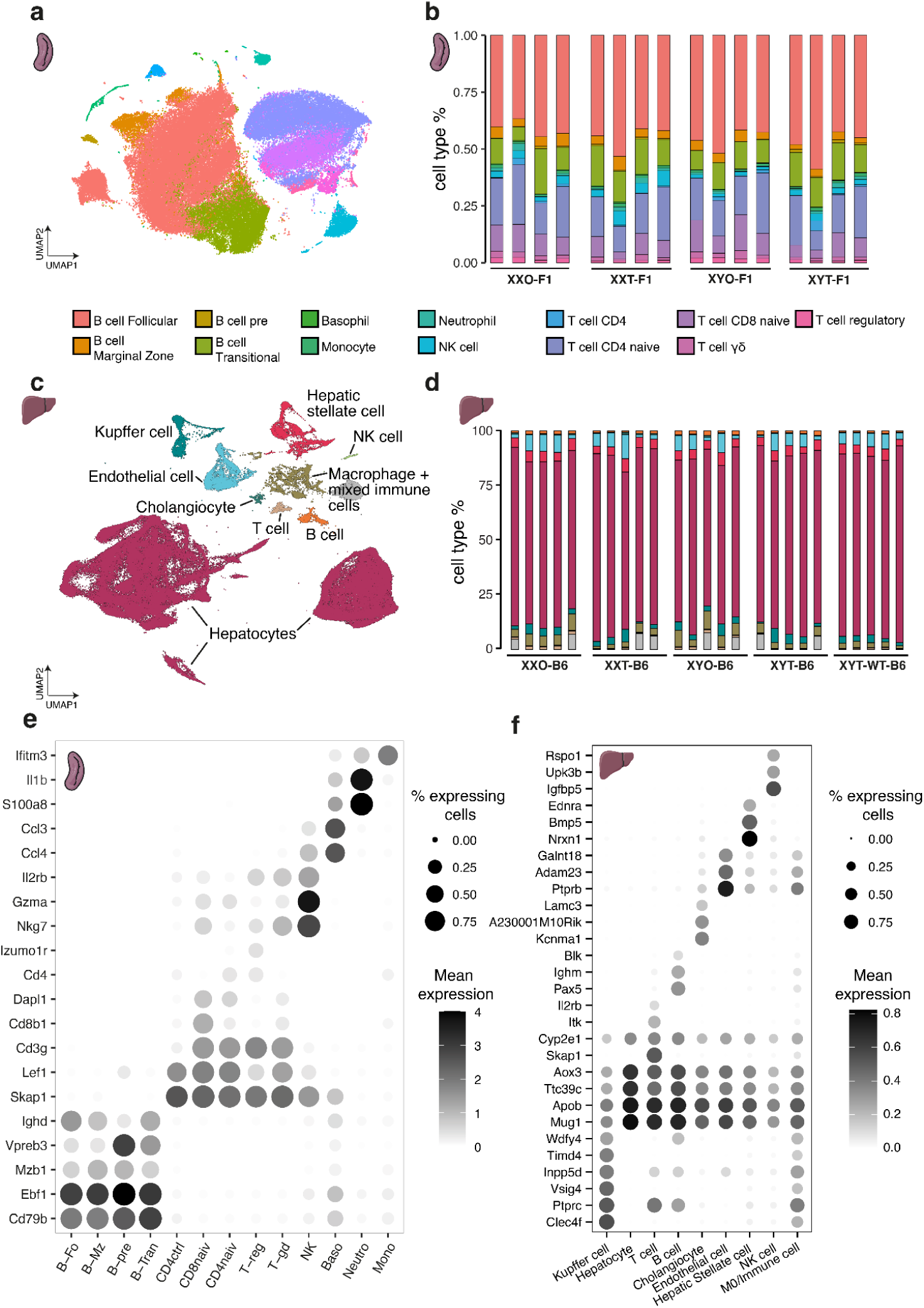
Single-cell dataset annotation. **a,c**, Uniform manifold approximation and projection (UMAP) plot of the sequenced cells. Cells are coloured based on cell type. **b,d,** Barplots showing % of each cell type for each biological replicate. **e,f,** Cluster and cell type-specific genes used to annotate cell types. For the spleen dataset, 103,000 cells were analysed with 13 unique cell type annotations. Consistent with previous reports, the splenic immune cell population of mice is dominated by lymphoid lineage cells. For the liver dataset, 147,888 cells from 25 individual mice were analyzed, with nine unique cell types identified.

**Supplementary Figure 4:**
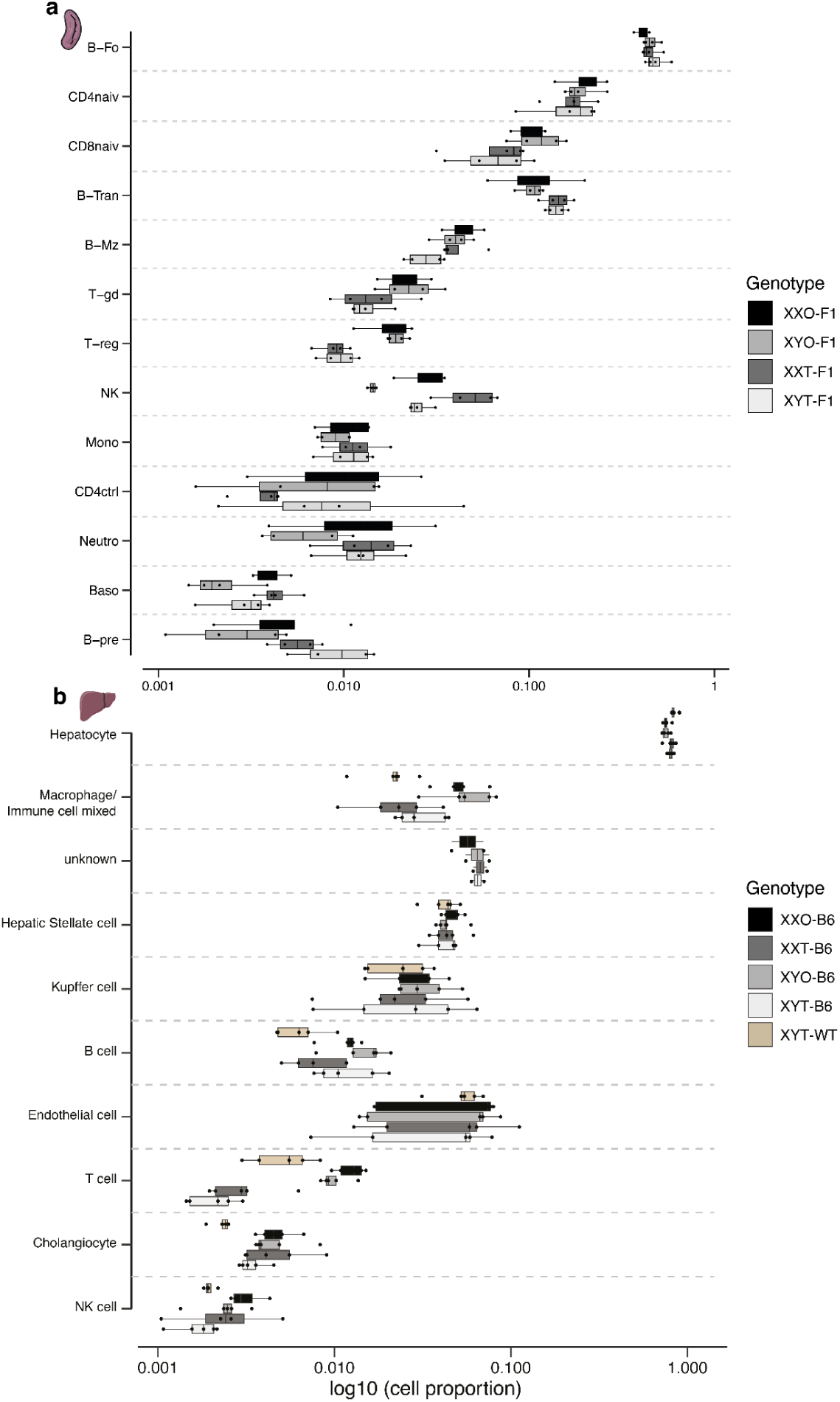
Few changes in cell type proportions across FCG genotypes. Cell type proportion comparison between the different genotypes in the spleen (**a**) and liver (**b**) dataset. Changes in cell type proportions between the FCG genotypes are small and mostly driven by differences in gonadal hormones.

**Supplementary Figure 5:**
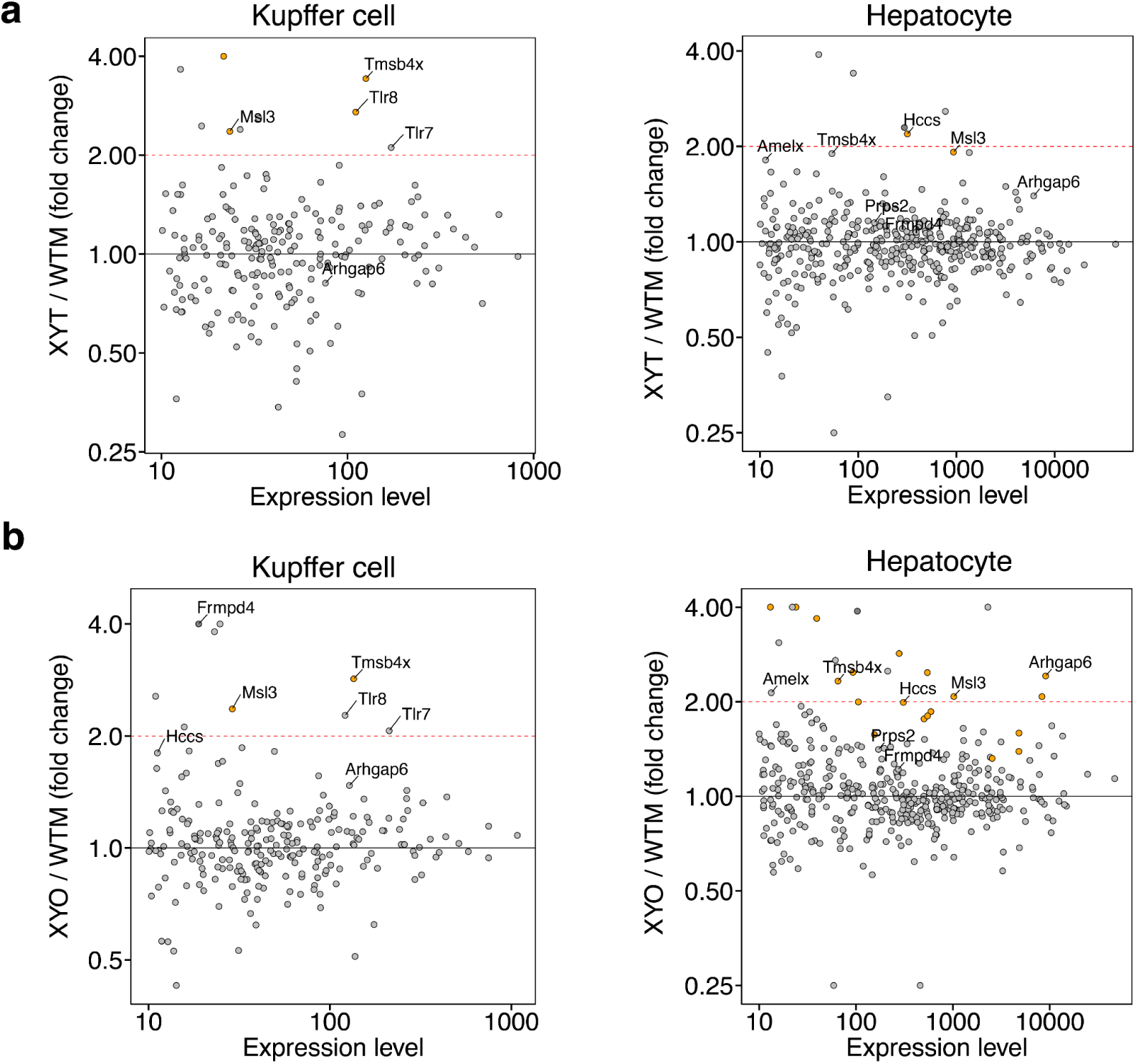
XY-FCG comparison to wild-type males. **a-d**, Differential expression analysis between FCG XY genotypes in pseudobulked Kupffer cells and Hepatocytes in the liver sci-RNA-seq data set.

**Supplementary Figure 6:**
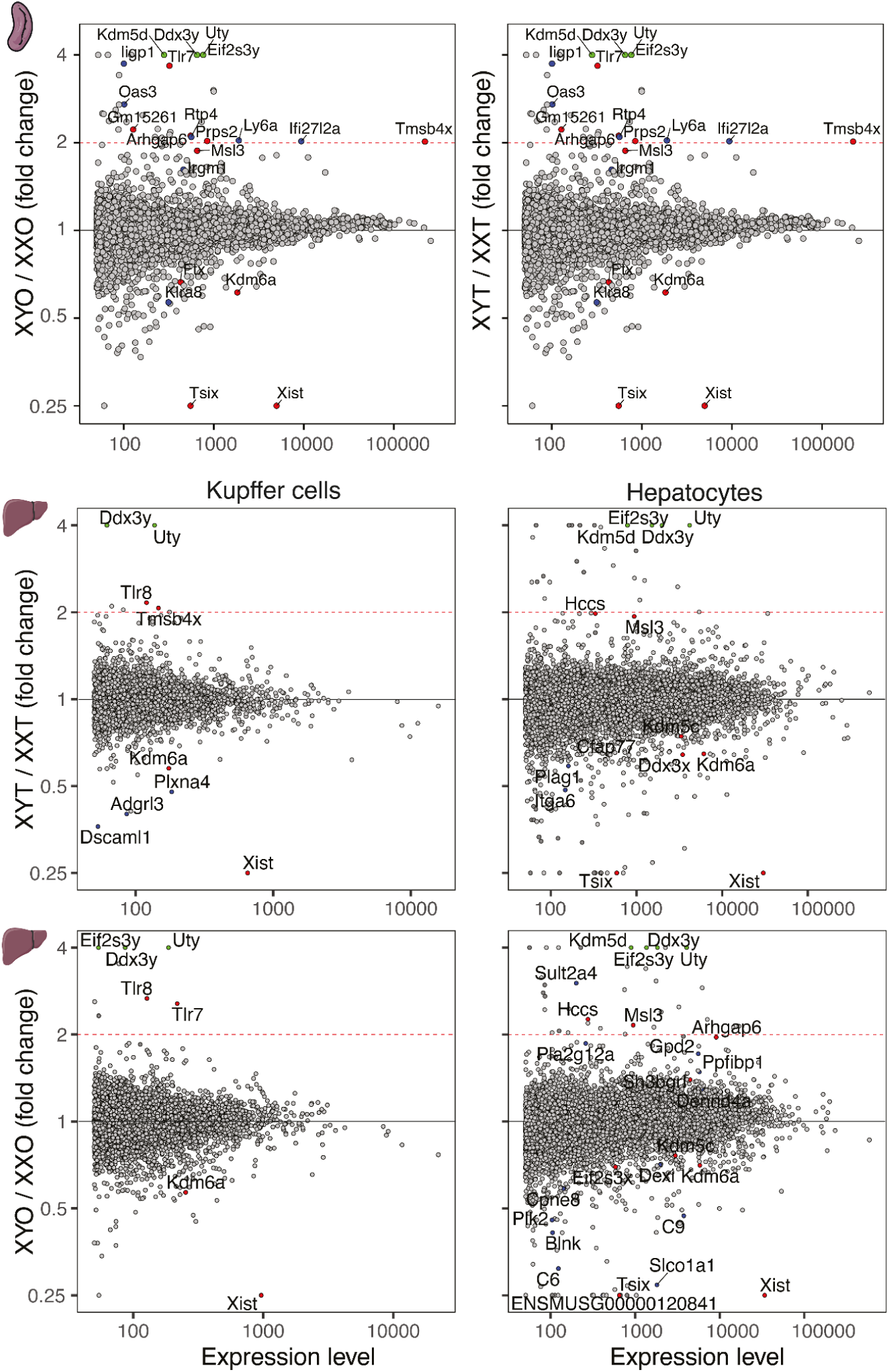
The increased expression of the duplicated genes has a minor effect on autosomal gene expression. Differential expression analysis of pseudo-bulked single-cell RNA sequencing from all spleen cells between XYT and XXT mice and between XYO and XXO mice. The coloured dots represent genes with a FC >=1 and padj>=0.1. Red dot: X chromosome genes. Green dot: Y chromosome genes. Blue dot: autosomal genes. Some autosomal genes, in particular *Ly6a* and *Ifi27l2a,* which we detect as differentially expressed in spleen in both comparisons are interferon target genes, potentially downstream of *Tlr7*.

**Supplementary Figure 7:**
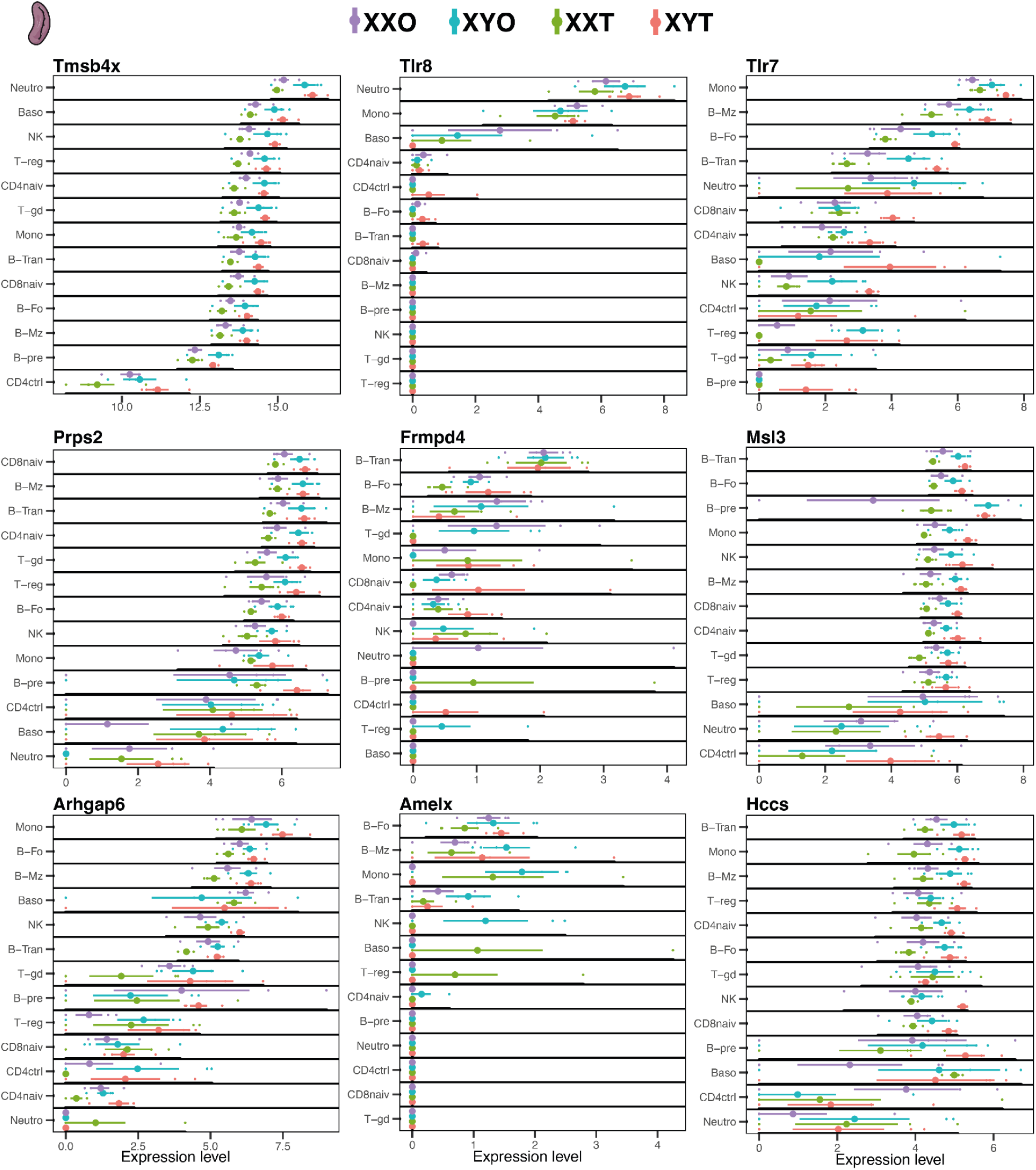
Expression of the duplicated genes in each of the genotypes in the spleen dataset. Independent measurements of all the biological replicates are shown as small dots. The main dot represents the mean of normalised expression with the standard error. The duplicated genes show cell-type specific expression and mostly a higher expression in FCG-XY genotypes.

**Supplementary Figure 8:**
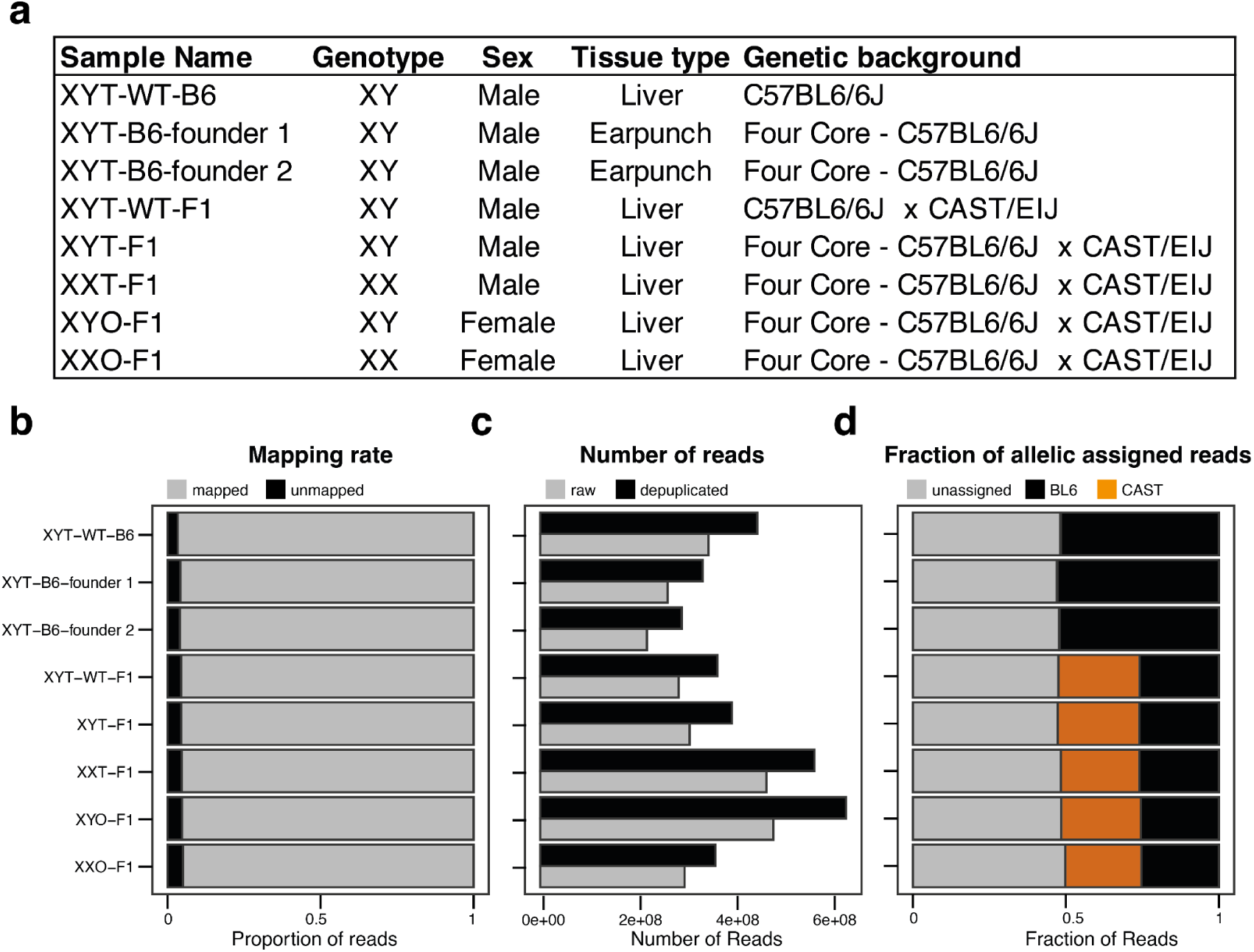
Quality control of the whole genome sequencing data. **a** Summary table of the samples used for whole-genome sequencing analysis. **b,** Barplots showing the mapping rate across samples. **c,** Barplots showing the number of mapped reads, before and after duplicate exclusion. **d,** Barplots showing the fraction of reads assignable to the C57BL6/6J and CAST/EiJ haplotypes based on known heterozygous single-nucleotide variants.

**Supplementary Figure 9:**
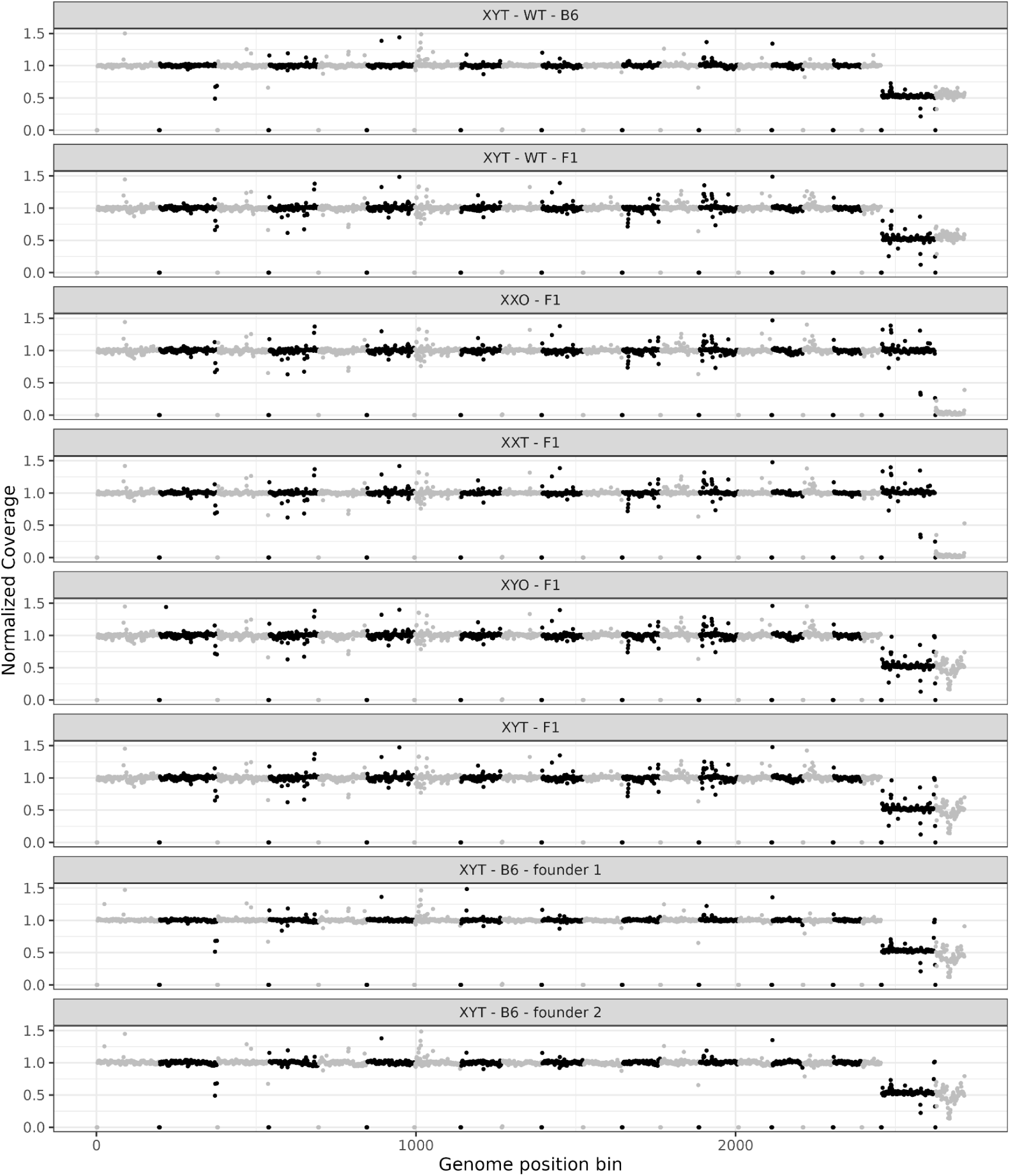
Genome-wide coverage plots across sequenced samples. Coverage is computed as the number of deduplicated, mapped reads in 1 mega-base windows, and normalized to the median across the sample, corresponding to a diploid genome. Chromosomes are colored alternately in black and grey, and the last two chromosomes are X (black) and Y (grey).

**Supplementary Figure 10:**
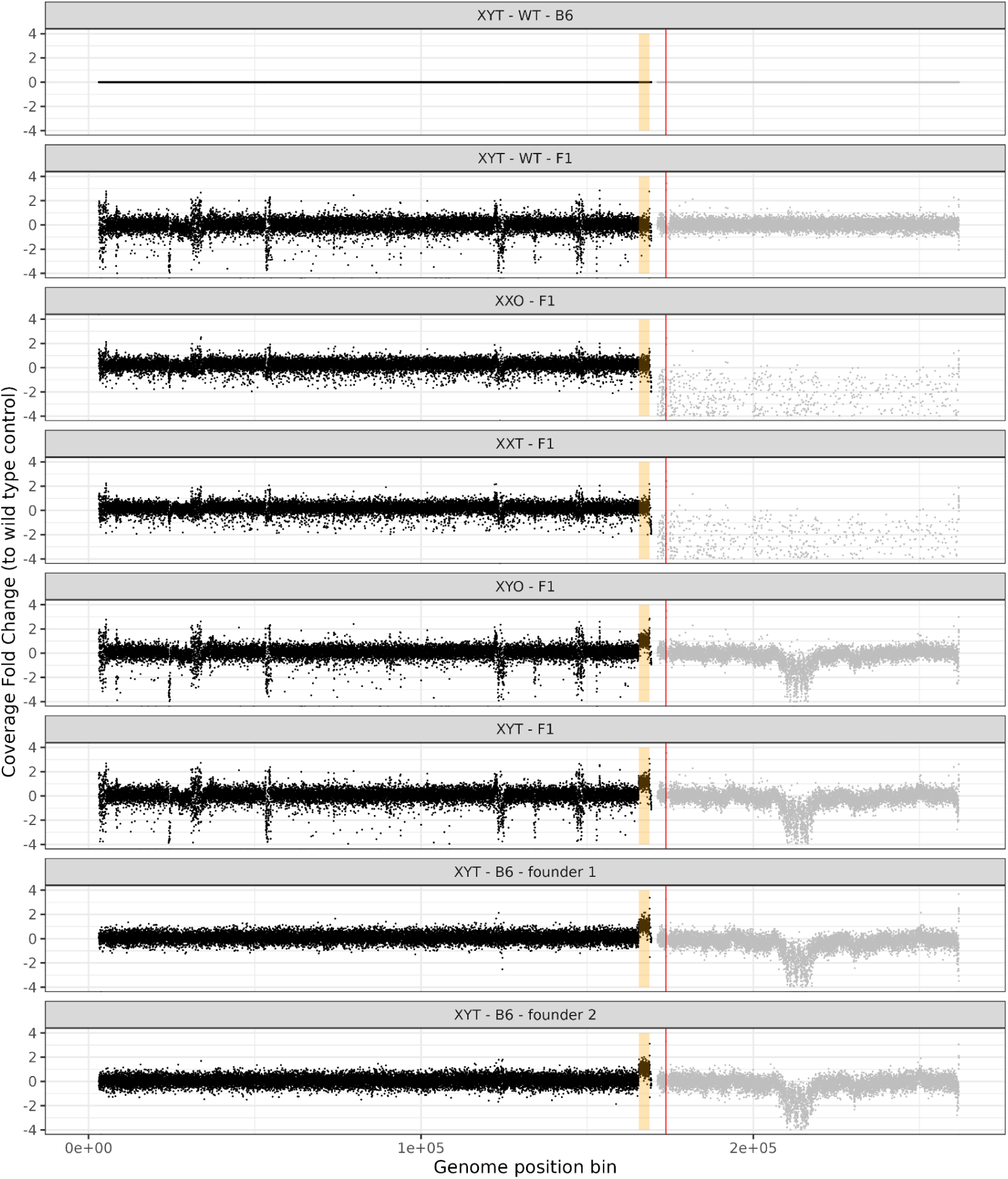
Sex-chromosome coverage plots across sequenced samples. Coverage is computed as the number of deduplicated, mapped reads in 1 mega-base windows, and normalized to the median across the sample, corresponding to a diploid genome. The X-chromosome is shown in black, the Y-chromosome in grey. In orange, the putative amplified region is highlighted. In red, the location of the *Sry* gene is shown.

**Supplementary Figure 11:**
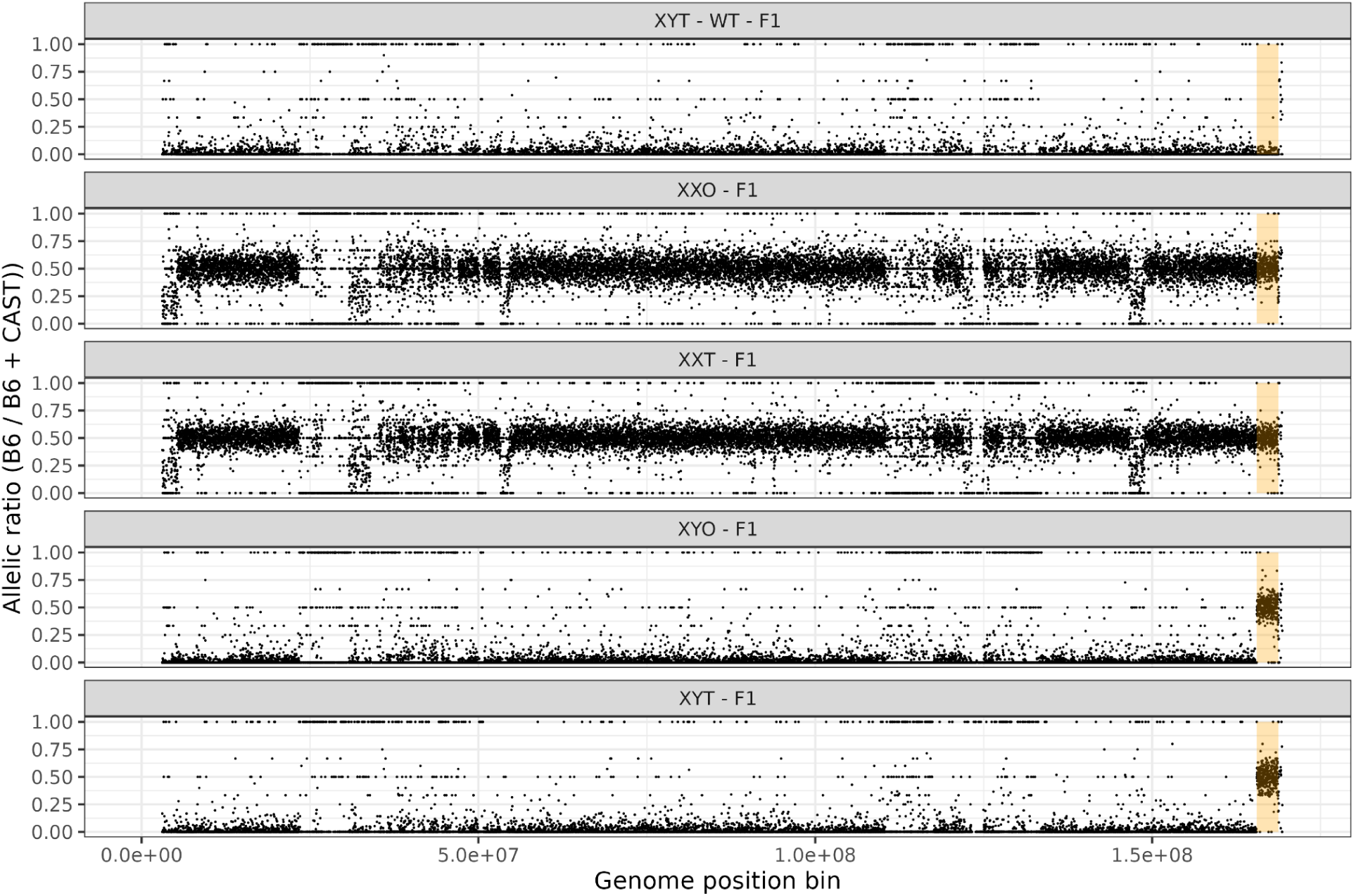
Allele-specific coverage plots on the X-chromosome across sequenced samples. Reads are assigned to paternal (B6) or maternal (CAST) haplotypes based on known homozygous variants between the two strains. Coverage per haplotype is computed as the number of deduplicated, mapped reads in 1 mega-base windows and allelic ratios B6 / (B6 + CAST) are computed. In orange, the putative amplified region is highlighted. As the X-chromosome is maternally inherited, the presence of B6 X-chromosomal reads in XY mice demonstrates a transfer of X-chromosomal material to a different chromosome.

## Notes

### Competing Interest Statement

The authors have declared no competing interest.

### Summary of Updates

Table 1 headers revised

